# Comprehensive Unbiased Analysis of Vascular Tissue Changes in Accelerated Atherosclerosis Using High-Resolution Ultrasound combined with Photoacoustic Imaging

**DOI:** 10.1101/2024.10.30.621032

**Authors:** Alwin de Jong, Valeria Grasso, Kayleigh van Dijk, Thijs. J. Sluiter, Paul. H.A. Quax, Jithin Jose, Margreet R. de Vries

## Abstract

Venous bypass grafts are commonly used to circumvent complex coronary or peripheral artery occlusions. The patency rates, however, are hampered due to accelerated buildup of atherosclerotic lesions in the vein graft wall. Identification of unstable plaques is crucial to guide clinical decision making. In this study, we employ advanced high-resolution ultrasound (US) coupled with spectral photoacoustic imaging (sPAI) to enhance the accurate visualization and analysis of tissue composition *in vivo*. By applying unbiased spectral analysis, we investigate the composition and plaque instability in a murine vein graft model.

**Method:** Male hypercholesterolemic ApoE3*Leiden mice and normocholesterolemic C57BL/6 mice underwent vein graft surgery in which a caval vein from a donor mouse was interpositioned into the arterial circulation of a recipient at the sight of the right common carotid artery. US imaging with sPAI was conducted on 7, 14, 21, and 28 days after surgery. Spectral curves from the near-infrared (NIR) I region, spanning 680 to 970nm, were extracted using a data-driven approach. Component discovery and cross-correlation analysis were performed with Matlab, and ImageJ reconstructed the components within 3D images. At the endpoint histological analysis of the vein grafts was performed.

**Results:** Analysis of the NIRI region revealed distinct components, with 7 and 10 components tested in the cross-correlation map. Relative abundance values identified melanin, oxidized hemoglobin, deoxygenized hemoglobin, lipids, and collagen. Lipids and collagen spectra accurately identified lipid and collagen-rich tissues *in vivo*. The sPAI analysis of of the vein graft wall *in vivo* resulted in a 8.7% lipids in the vein graft wall compared to 1.8% lipids in the histological analysis at t=28d. For vein grafts from ApoE*3-Leiden mice no differences in the lipid positive area was observed between the sPAI analysis or histological quantification. The percentages collagen present in the vein graft walls from both strains analyzed via sPAI and histological showed comparable results at t=28d.

**Conclusion:** Our study demonstrates that sPAI can be utilized for compositional analysis of murine tissue in an unbiased manner. This methodology can be used to enhance our understanding of vein graft dynamics and holds promise to advance non-invasive characterization of vascular diseases to ultimately guide clinical decision making.

## 1. Introduction

Defining tissue composition is crucial to enhance early disease detection and monitor disease progression. Photoacoustic imaging (PAI) is a hybrid imaging modality that uses the optical absorption properties of tissue chromophores to obtain compositional information which is combined with the spatial resolution of ultrasound, thus offering a non-ionizing (and non-invasive) method for detailed tissue analysis. In PAI, pulsed laser lights are delivered into tissue, which absorp the optical energy. Consequently, this causes thermoelastic expansion of the tissue, generating acoustic signals that can be detected via ultrasound. Due to the unique emission spectrum of every molecule, various tissue components can be distinguished when multiple wavelengths are measured. In vivo PAI can be conducted in the near infrared region-1 (NIR1) and –2 (NIR2), ranging from 700-900 nm and 1000-1700 nm respectively. Currently, NIR1 is most frequently used for PAI, although NIR2 offers greater penetration depth and decreased auto-fluorescence. Unfortunately, NIR2 imaging is hampered by low efficiency above 1100nm, thus significantly impacting tissue analysis.

Molecular imaging can enhance early disease detection and aid in monitoring disease progression through visualization and quantification of tissue components. Photoacoustic imaging (PAI) is a hybrid imaging modality that combines the optical absorption properties of tissue chromophores with the spatial resolution of ultrasound, offering a non-ionizing method for detailed tissue analysis^1, 2^. PAI employs pulsed laser light to excite tissue, which absorbs the optical energy. This causes thermoelastic expansion of the tissue, generating acoustic signals that can be detected via ultrasound^3^. The emission of acoustic signals depends on the optical absorption of individual tissue components. When multiple wavelengths are measured, this allows for identification of different molecular components in tissue^4^.

Recently, spectroscopic photoacoustic imaging (sPAI) has shown promise in the quantification of individual tissue components in various disease models, including tumor microenvironments, hemoglobin content, and atherosclerosis^5-8^. Due to strong hemoglobin absorption below 600 nm and challenges posed by water absorption and decreased imaging efficiency above 1100 nm, in vivo PAI is typically conducted within the 600–1100 nm range. ^9^. However, imaging in the near-infrared region (NIR) byond 1100nm offers deeper tissue penetration and improved specificity^10, 11^.

Identification of molecules often occurs via linear unmixing, which requires the input (spectral curves) of expected tissue components. This is challenging since the known spectra might differ from the actual spectra that are detected in disease conditions. Moreover, the biased approach hinders identifications of novel components. To overcome this issue, we have recently developed an unsupervised superpixel photoacoustic unmixing (SPAX) framework that enables fully automatic spectral unmixing and accurate quantification of molecular components. During atherogenesis, accumulation of lipids in the vessel wall leads to excessive infiltration of immune cells, which together fuel continuous growth of the plaque. In due time, atherosclerotic plaques can become unstable and rupture, leading to e.g. myocardial infarction or stroke. Currently, there are no options to obtain compositional information on atherosclerotic plaques, which can potentially be used to identify high-risk plaque that warrant invasive vascular interventions.

In the current manuscript, we therefore investigated the performance of the SPAX framework to analyze the composition of atherosclerotic blood vessels. To this end, normo– and hyper-cholesterolemic mice underwent venous bypass surgery. Over time, the wall of the venous conduits thickens, whilst hypercholesterolemia also triggers buildup of lipids. Using PAI and the SPAX framework, we are able to identify multiple components in the vessel wall such as (oxygenated) hemoglobin, lipids and collagen. Lipids and collagen deposition in the vein graft wall corrolate with disease severity^12^ and is a valuable parameter in vein graft remodeling^13^. Lipids accumulate under hypercholesterolemic conditions in the vein graft wall, that can be identified over time supporting the histoligcal analysis post-mortem.

## 2. Materials and methods

### 2.0. Study approval and mice

This study was performed in compliance with the Dutch government guidelines and the Directive 2010/63/EU of the European Parliament. All the experiments were approved by the institutional committee for animal welfare of the Leiden University Medical Center. Male C57BL/6 and C57BL/6 ApoE*3-Leiden^+/-^ (ApoE*3-Leiden) mice were fed a western type diet containing 1.0% cholesterol and 0.5% cholate (HFD) (Sniff Spezialdiäten, GMBH, Soest, Germany) from 8 weeks of age. The mice were bred in our own institute and had free access to food and water for the entire duration of all experiments. Heterozygosity of *ApoE*3-Leiden* results in hypercholesterolemia of the *ApoE*3-Leiden* mice, whilst the wildtype C57BL/6 mice have normal serum cholesterol levels.

### 2.1. Phantom analysis of collagen and lipids *in vivo*

Spectral mapping of collagen type I, (Rat tail, #08-115 Merck) and gelatin (Type I and III, #G2500, Merck) was performed using the Vevo LAZRX Phantom analysis. Collagen and gelatin was loaded on 0.16mm^2^ tubing (FUJIFILM Visual Sonics) in water to facilitatie image aquisition. Spectral imaging was acquired in the NIR I region 680nm – 960nm with 5nm steps. For lipids, the inguinal fat pad was used to for spectral mapping in the NIR I region. For brown adipose tissie, the cervical-thoracic region has been imaged *in vivo*, and *ex vivo*. After these acquisitions, the interscapular fat pad from the cervical-thoracic area was dissected post-mortem and used for spectral measurements. Spectral datasets were analysed with Matlab r2023b.

### 2.2. Vein graft surgery

After 28 days of HFD, mice were anesthetized with an intraperitoneal (i.p.) injection consisting of a combination of midazolam (5 mg/kg, Roche), medetomidine (0.5 mg/kg, Orion), buprenorphine (0.1 mg/kg, MSD Animal Health) and fentanyl (0.05 mg/kg, Janssen). Adequacy of anaesthesia was monitored by toe pinching. Caval veins were obtained from surplus C57BL/6 mice, which were also anesthesized with the aforementioned combination of drugs. These caval veins placed into the arterial circulation of recipient C57BL/6 and ApoE*3-Leiden mice at the site of the right common carotid artery, as previously described^14^.The morning after surgery and on indication, additional buprenorphine was given for pain relief. 28 days after surgery, these mice were anesthetized, exsanguinated via blood draw through the orbital sinus after which the thorax was opened and the vasculature was perfused with phosphate-buffered saline (PBS, Braun) from the left ventricle. Thereafter, vein grafts were harvested, snap frozen and stored at - 80 °C.

### 2.3. Ultrasound and sPAI

For PAI, animals were anesthetized with isoflurane and placed on the animal imaging platform of the Vevo LAZR-X system, where temperature, heart rate, and respiration rate were monitored in real-time. During the experiments, anesthesia was maintained using a vaporized isoflurane (1 L/min of oxygen 0.3L/min air, and 2.5% isoflurane) gas system. The mouse was positioned in right lateral recumbency, and the transducer was aligned perpendicularly. High-resolution Ultrasound (US) and Photoacoustic (PA) imaging (US-PA) have been acquired by using the platform Vevo LAZR-X (FUJIFILM VisualSonics, Inc., Toronto, ON, Canada). A linear US transducer array (MX550S) consisting of 256 elements at a nominal center frequency of 40 MHz and bandwidth of 25-55 MHz, is coupled with narrow optical fibers, mounted on either side of the transducer. This high-frequency transducer guarantees high spatial resolution (axial resolution of ≈40⍰µm and lateral resolution of ≈80⍰µm), and the narrow optical fibers enable focused illumination. Homogenous light illumination is guaranteed by placing the sample to be imaged on the converging area of the two light beams. Spectral photoacoustic imaging (sPAI) was performed on the short axis at three separate regions of the vein graft within the wavelength range of 680-970 nm with a 1nm step. Imaging acquisition was conducted on days 7, 14 and 28 after surgery. Additionally, to visualize the entire vein graft, 3D multiwavelength photoacoustic images were acquired at 680, 700, 720, 740, 760, 780, 800, 820, 840, 860, 880, 900, 910, 920, 930, and 960nm. During the volumetric multi-spectral PA acquisitions, a stepper motor is used for the linear translation of the US transducer and optical fibers along the sample. The linear stepper motor moves in steps of 0.08mm increments while capturing 2D parallel images, for a 3D range distance of 4*mm*. Image visualization, reconstruction, and processing were realized with VevoLAB 3.2.6 software (FUJIFILM, VisualSonics)^15^.

### 2.4. Spectral superpixel unmixing analysis

A newly developed superpixel data-driven unmixing (SPAX) framework described in detail elsewhere^16^ was used for the blind tissue composition analysis. In addition to the automated detection of the oxidized hemoglobin and total hemoglobin (Hb) content, the spectra of lipid and collagen as well as their spatial distribution was also obtained by the unsupervised SPAX approach. The framework models the sPAI as a linear mixture M=WH, where abundance maps W and source spectra H are both unknown. The SPAX implements an SVD-based analysis to automatically distinguish the relevant spectral information above the noise level. SPAX is also extended to compensate for the spectral coloring artifact combining US image segmentation and spectral Monte Carlo (MC) light fluence simulations based on a predefined library of tissue optical properties. Besides, the advanced superpixel subsampling integrated within SPAX enables to detect less and most prominent components, without a priori information. After the spectral unmixing, the identified endogenous tissue chromophore spectra were correlated with the theoretical absorption spectra^17^ The longitudinal correlation between the extracted oxydized/deoxyzided hemoglobin, lipids, and collagen spectra and the theoretical absorption spectra has been evaluated. The correlation can give an indication of spectral changes that might be caused by inflammatory responses.

On the other end, from the normalized unmixed maps of lipid and collagen, obtained as output of the SPAX analysis, and a manual selection of ROI in the vein graft cross section we have evaluated the presence of the component within the selected area. This ensures a more accurate evaluation, since eventual artifact pixels coming from the complete FOV are excluded. The SPAX processing of the spectral 2D and 3D PA data is performed offline, after the acquisition. The computational time ranges between 1–12 min, depending on the size of the input sPAI data and the computer specifications.

### 2.5. Morphometric and compositional analysis

The vein grafts were cryo-sectioned in sequential cross-sections of 5µm thick made throughout the embedded vein grafts. The total vein was analyzed by minimum of 6 equally spaced sections. The plastic cuff served as the starting point for mounting sections onto glass slides. From this staining, the lumen areas, intimal areas, and total vessel areas were measured. Sirius red staining (Klinipath 80,115) was used to quantify the amount of collagen I/III present. Lipids were visualized with the oil-red-o staining (Abcam, ab150678). Slides were scanned using a slide scanner (3DHistec) and analyzed with ImageJ (FIJI). The measured area positive for either collagen or lipids was normalized to the size of the vein graft wall, yielding % positive area.

### 2.6. Statistical analysis

Differences in continuous variables between groups were statistically assessed by using the unpaired parametric t-test in Graph Pad Prism software(10.2.3). Data are represented as means ± SD unless stated otherwise. Significance was set at P<0.05. Significant differences are graphically represented as * P<0.05, ** P<0.01, and *** P<0.001.

## 3. Results

### 3.0. Generation of a spectral library to identify vein graft tissue components

To identify different spectral components of the vein graft wall, a spectral library was generated from vein grafts of ApoE*3-Leiden mice (n=12) at day 28 after engraftment (Fig. 1A). Color doppler signal was used to discen the vein grafts (Fig. 1B) and determine the region of interest in the PA images accordingly (Fig. 1C). Spectral curves (wavelength range 680-970nm, 1nm stepsize) were obtained from the vein grafts (Fig. 1D) on the short-axis view at three different locations (Fig. 1E). From these spectral images, the SPAX algorithm identified ten unique components (Fig. 1F).

**Figure 1.**
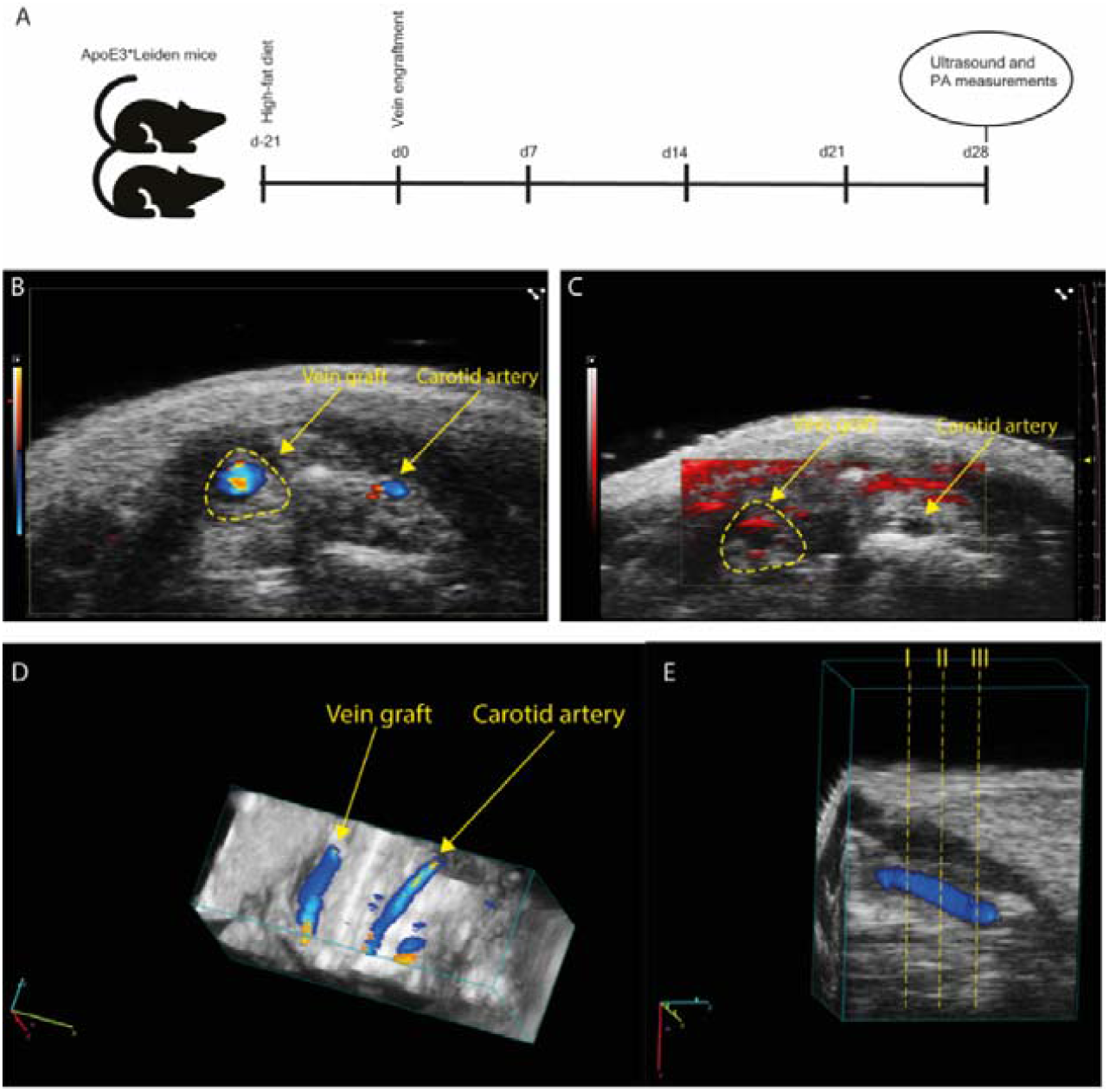
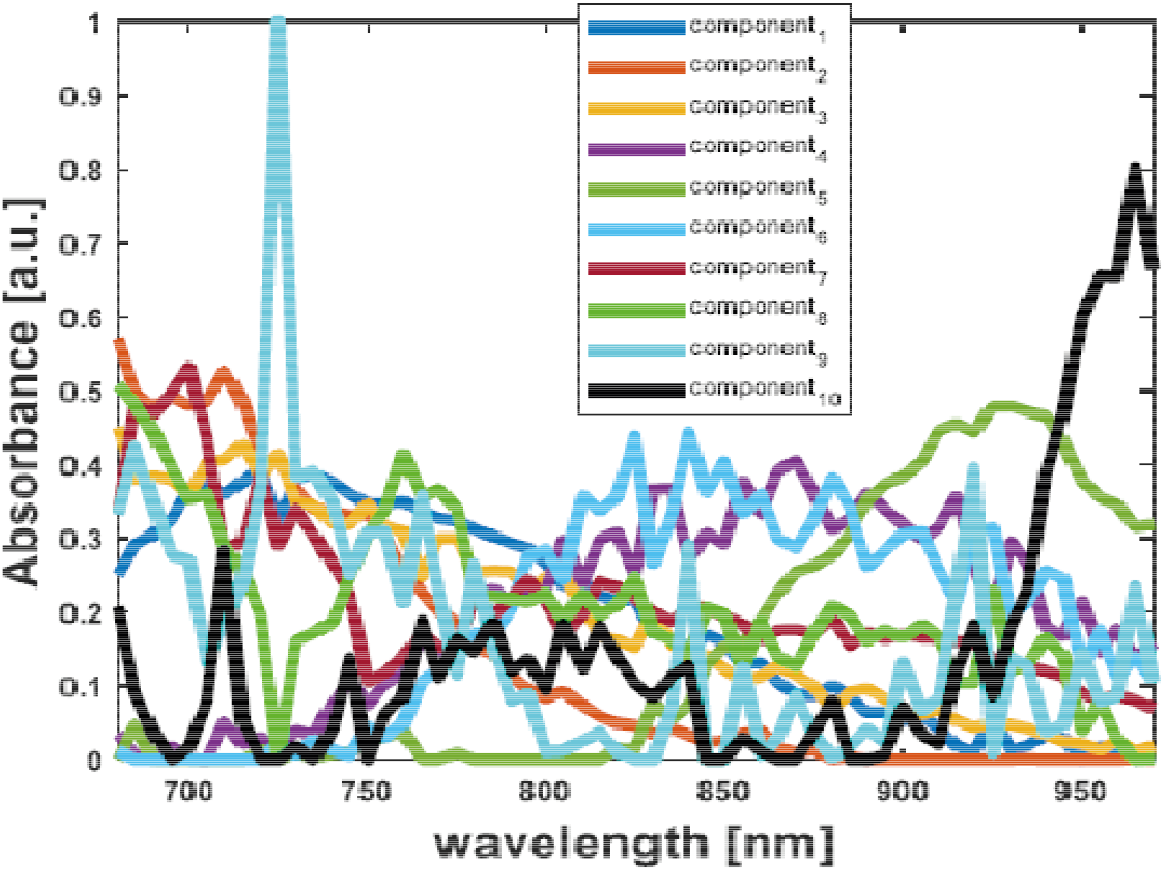
Vein graft PA acquisition and NIR1/NIR2 spectral curve identification. ApoE*3-Leiden mice were fed a high-cholesterol diet prior to vein engraftment, and spectral imaging was performed at t=28d **(A)**. The vein graft was visualized by color Doppler imaging **(B)** and spectral imaging **(C)**, with the vein graft highlighted by the yellow dotted line. From a 3D scan **(D)**, three different regions were selected for spectral imaging in the NIR region **(E)**. From these spectra, the SPAX algorithm isolated ten different components **(F)**.

#### Spectral cross correlation analysis identify 6 unique spectra in the vein graft wall

Within the ten different spectral curves that were identified, six correlated with known spectra that have previously been described in literature and (Fig. 2A). Windowing increased the differences in spectra that are more different and increased the alikeness in spectra that were more similar to reduce noise and artifacts in the spectral library (Fig. S1).

**Figure 2.**
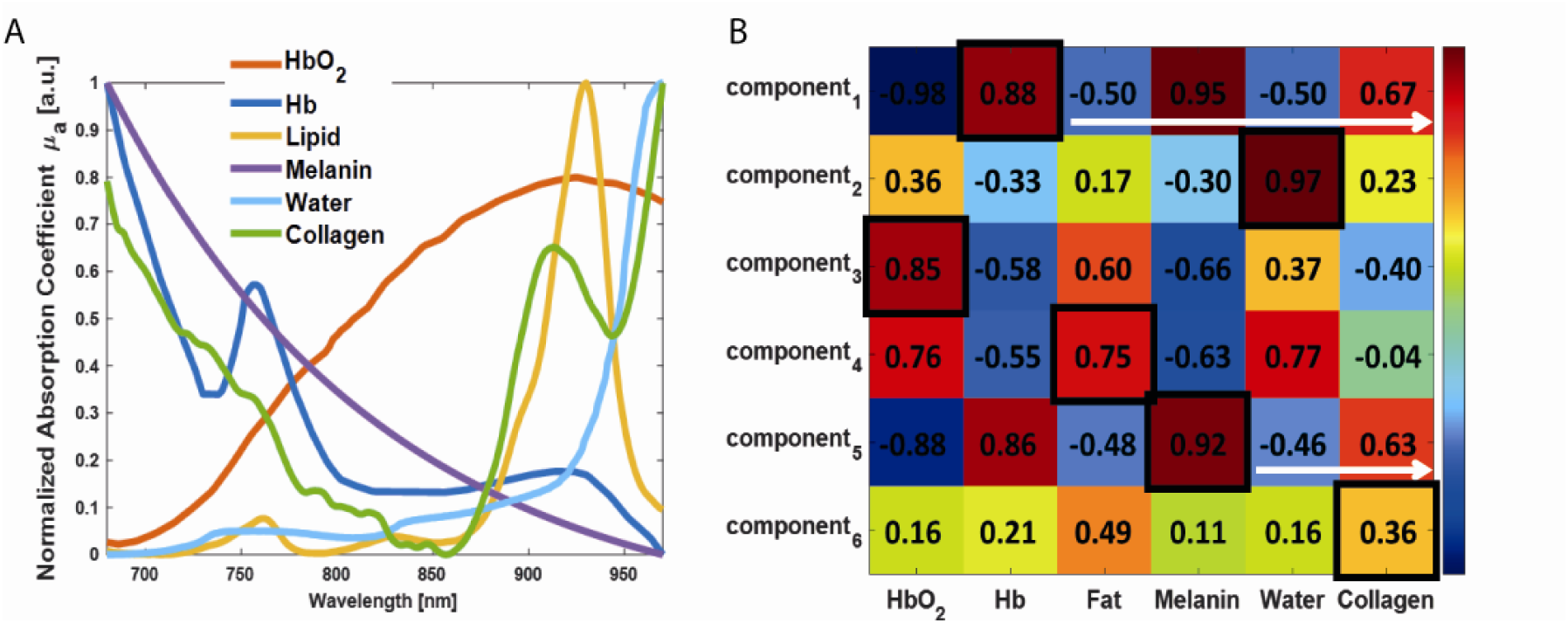
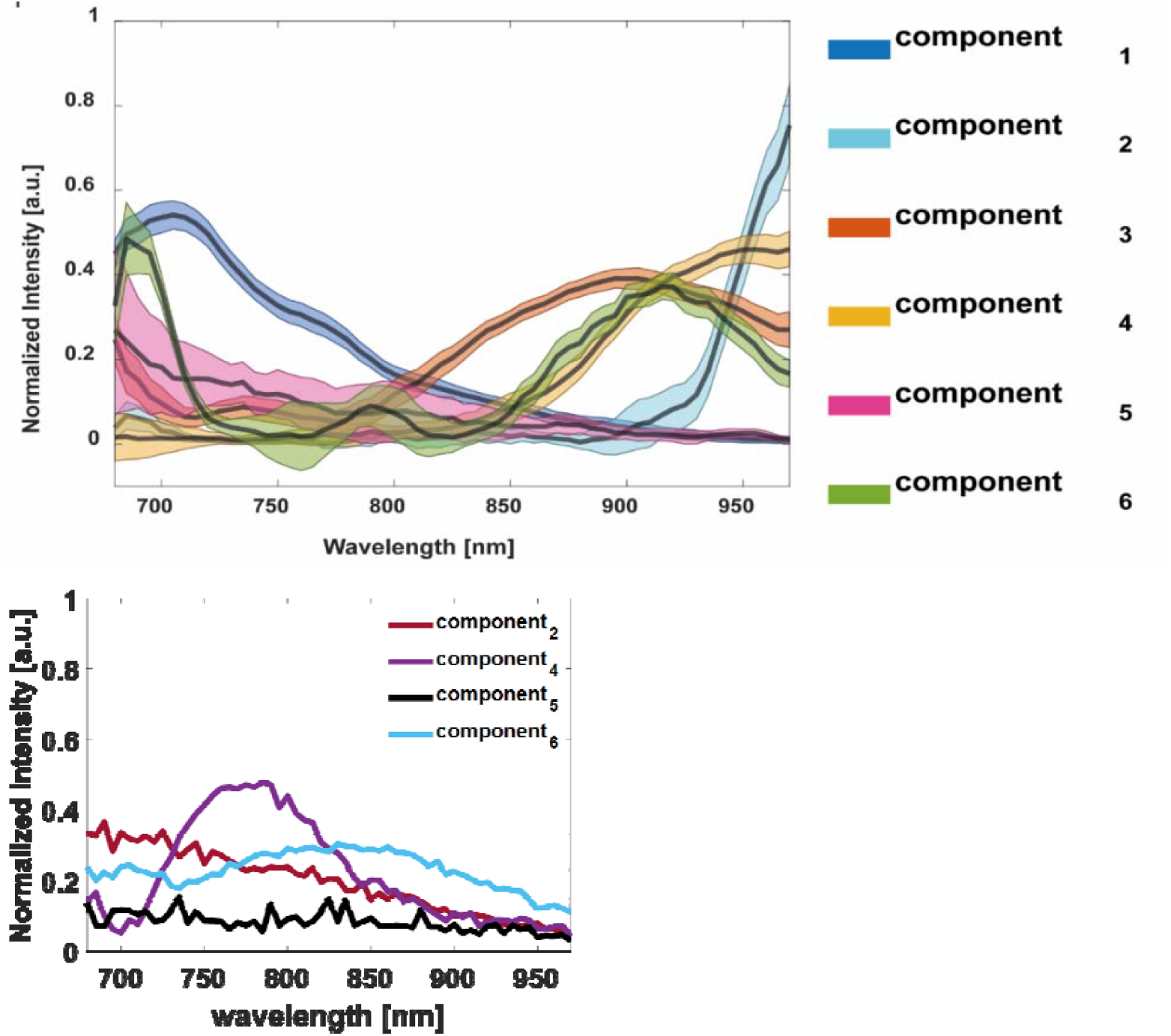
Vein graft PA acquisition and NIR1/NIR2 spectral curve identification. The known spectra of oxygenated hemoglobin, deoxygenated hemoglobin, lipids, melanin, water, and collagen are shown **(A)**, along with the cross-correlation of the known spectra with the identified spectra from the t=28d spectral data. The components were cross-validated with known spectra of different vascular tissues, resulting in a cross-correlation table **(B)**. A value close to 1 indicates spectra that are more similar, while a value closer to 0 indicates spectra that are different. A negative value means that the components are mirrored compared to the known spectrum. Indicating that spectrum 1: hemoglobin, 2: water, 3: oxidized hemoglobin, 4: lipids, 5:melanin, 6: collagen **(C)**. Four spectral curves were identified but could not be referenced against known spectra **(D)**.

As part of the extracellular matrix, lipids and collagen are present in the vein graft wall. Collagen is a contributor to vein graft wall stability while lipids are drivers of atherosclerotic lesion progression ^18^. The visualization of collagen and lipids, are identified as components four and component six, with a correlation of 0.75 and 0.36 with the reported spectra respectively (Fig. 2B). From the ten different spectra, six could be classified as tissue components present in the vein graft wall. However, the other four spectra that were identified could not be classified because reference spectra have not been found. One of these spectra display a peak at 760 – 785nm, while the other three shows a gradual decrease in intensity classified as a non-specific signal (Fig. S1). Although the correlation coefficient is >0,5 the relative abundance is more important in the validation of the isolated spectra because it provides a measure of how much of each component is present in the sample. Even if the spectral signature of a component matches a known reference spectrum with a high correlation coefficient, it may not accurately represent the composition of the sample if its abundance is low. This shows that the identified components can be unbiased classified and act as a spectral library. The data-driven SPAX analysis has been applied longitudinally to the spectral data of ApoE*3-Leiden mice (n=12) used as reference group. From the automated unmixing six source spectra have been identified (see Fig. 1F). Fig. 1F depicts the mean μ (solid lines) and standard deviation σ (shaded area) of the identified source spectra for ApoE*3-Leiden mice reference group.

The absortion spectra of endogenous tissue component obtained from literature^17^ and used as theoretical library are shown in figure 1G. These components are used for cross-correlation analysis with the automatically identified spectra via SPAX, to identify the extracted components. Fig. 2 (B) shows the correlation map obtained between the spectra of the theoretical library and the clustered spectra obtained from the SPAX analysis of the ApoE*3-Leiden mice (n=12) used as reference. Fig. 2B shows the cross-correlation map between the blindly unmixed components via SPAX and the library of spectra used as reference (see Fig. 2A). Specifically, the six unmixed source components obtained via SPAX have been identified via cross-correlation as: Hb (component_1), water (component_2), HbO2 (component_3), adipose tissue (component_4), melanin (component_5), and collagen (component_6) with pearson correlation coefficients of 0.88, 0.97, 0.85, 0.75, 0.92, and 0.36 respectively.

### 3.1. Lipid and collagen spectra validation in vivo

To substantiate the lipid and collagen spectra obtained from the in vivo vein graft spectral library, we also obtained spectral curves from interscapular brown adipose tissue (iBAT) in vivo as well as ex vivo. The spectral imaging of the BAT in vivo revealed a maximum intensity at 925 nm, a decrease at 935 nm, and a gradual increase until 970 nm, corresponding with the lipid reference spectral curve (Fig. 3A). The ex vivo and dissected BAT spectral curves shifted, reaching maximum intensity at 935 nm and gradually increasing until 970 nm (Fig. 2). Next, we evaluated whether sPAI could discriminate between BAT and white adipose tissue (WAT) in vivo. The spectral data from the inguinal fat pad peaked at 945 nm, without the gradual increase in intensity observed in BAT. This suggests that sPAI can differentiate between BAT and WAT in vivo. Next, we validated whether component_6 corresponded with collagen. Phantom analysis of collagen I/III revealed a gradual decrease from 680 nm to 970 nm, while gelatin, mainly composed of collagen I, showed a maximum intensity at 805 nm (Fig. 3B). Spectral scanning of the collagen patch isolated from the mouse tail resulted in a spectral curve with distinct peaks at 880 nm and 940 nm (Fig. 3B). This shows that component 6 from the spectral library is able to identify collagen in the sPAI dataset and component 4 shows lipids, with a difference in spectral signature between white and brown adipose tissue.

**Figure 3.**
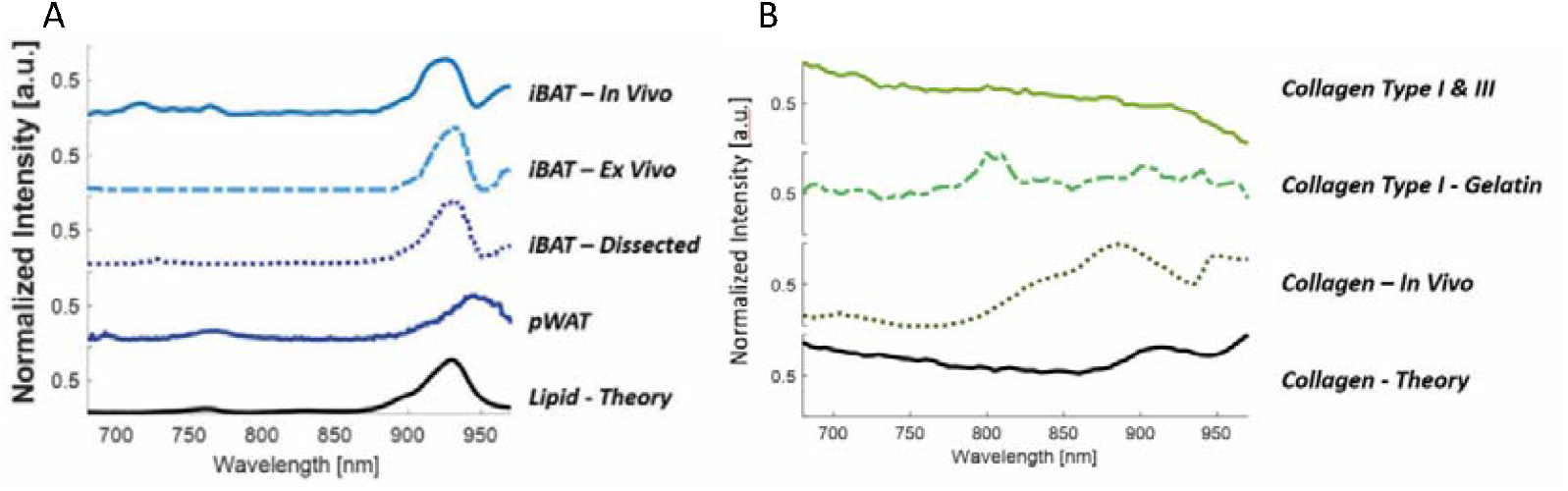
Validation of the white adipose tissue, brown adipose tissue and different types of collagen in vivo and PHANTOM analysis. Spectral analysis the intrascapular brown adipose tissue region of a BALB/c mouse, brown adipose tissue dissected **(A)**. The peripheral white inguinal fat pad is (pWAT) used as the imput for the white adipose tissue spectro resulting in different normalized intensities. PHANTOM analysis of collagen type I and III, gelatin, collagen in vivo from the tail of a rat and the theoretical spectrum of collagen **(B)**.

**Figure 4.**
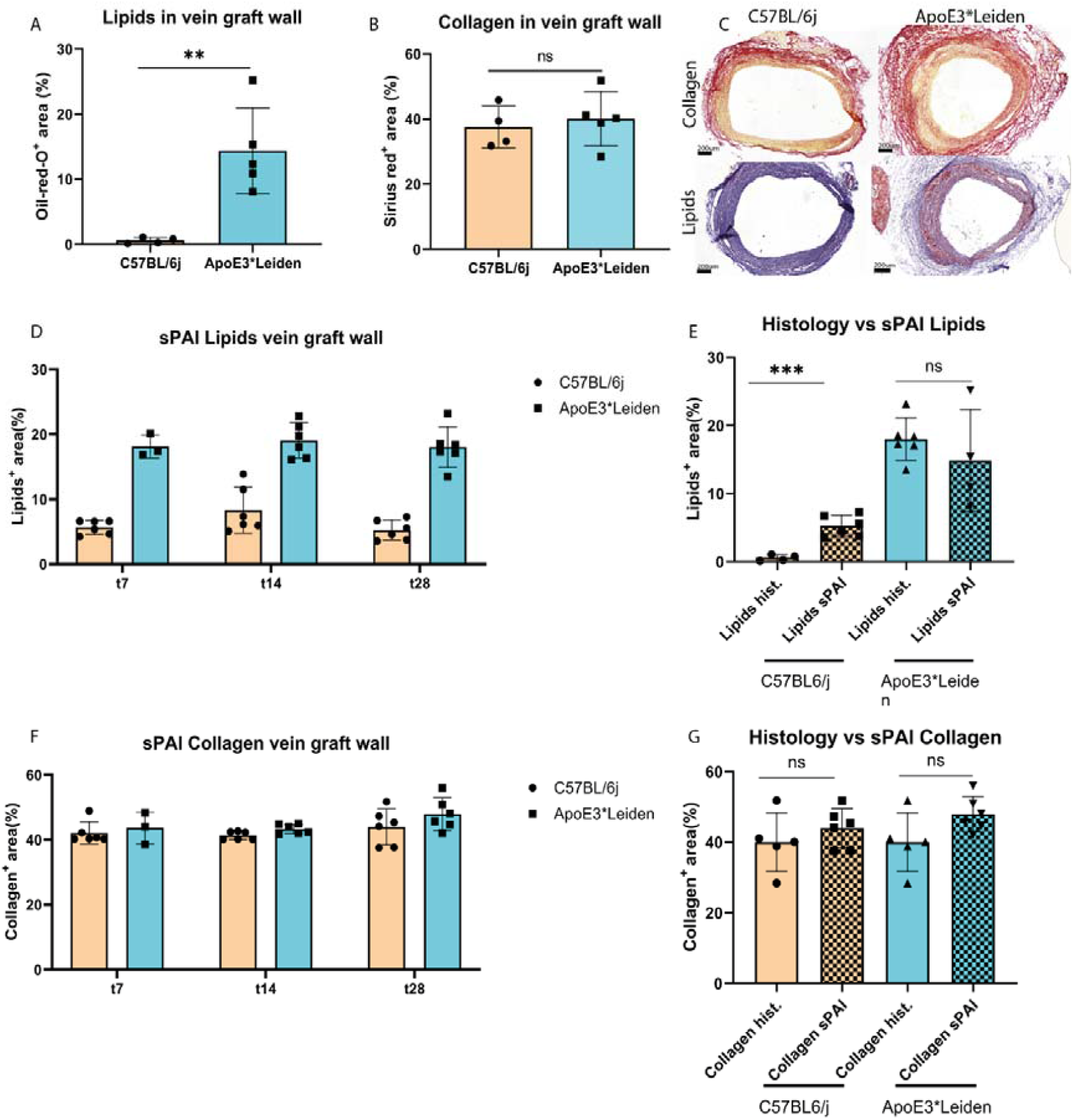
Non invasive compositional analysis of vascular remodeling over time, morphometrical validated on frozen tissue sections. After sacrifice, the vein grafts were snap frozen and the lipids and collagen quantified in the vein graft wall. sPAI analysis over time compared to histological analysis for the amount of lipids and collagen in the vein graft wall at day 28. Significant differences are graphically represented as * P<0.05, ** P<0.01, and *** P<0.001.

**Figure 4.**
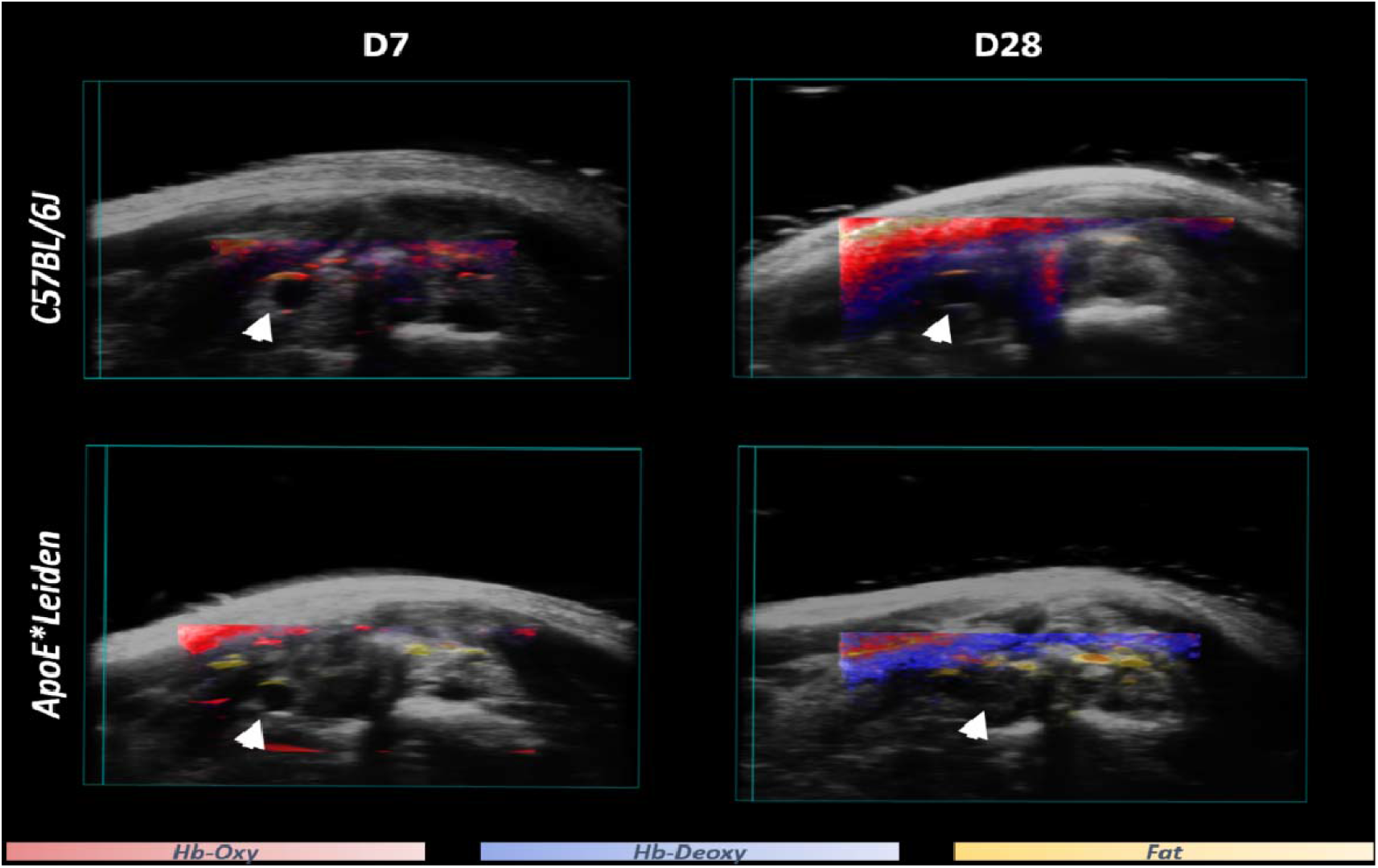
Location of different components within the 3D in vivo architecture. Visualisation of the vein graft (arrows) with the lipids and oxygenated / deoxygenated hemoglobin in ApoE*3-Leiden mice and C57BL/6 mice at day 7 and day 28.

### 3.2. Non invasive compositional analysis of vascular remodeling over time, morphometrical validated on frozen tissue sections

To longitudinally assess changes in vein graft composition in vivo, normo-cholesterolemic C57BL/6 and hyper-cholesterolemic ApoE*3-Leiden mice (n=4-5 / group) underwent vein graft surgery. 7, 14 and 28 days after surgery, 3D multiwavelength PA images were obtained. Unmixing of the data using the SPAX algorithm, with the 6 earlier identified spectra serving as baseline spectra, indicated that the lipid content of the vein graft did not change over time for the C57BL/6 as well as the ApoE*3-Leiden mice. Moreover, the lipid content was significantly increased in the ApoE*3-Leiden mice compared to C57BL/6 (p=0.0004). Histological analysis of vein grafts harvested at postoperative day 28 using oil-red-o staining confirmed buildup of lipids in the vein graft wall of ApoE*3-Leiden but not C57BL/6 mice. Additionally, analysis of the PA images demonstrated that collagen content was unaffected by mouse strain and did not change over time. Histological analysis using Sirius Red staining also revealed similar collagen content between C57BL/6 and ApoE*3-Leiden mice. The vein graft wall from ApoE*3-Leiden mice composed of 14.2% of lipids compared to 1.8% lipids in vein grafts isolated from C57BL/6 mice (P=0.008) while the amount of collagen present in the vein graft walls were not different between both strains of mice. The sPAI analysis of of the vein graft wall *in vivo* resulted in a 8.7% lipids in the vein graft wall compared to 1.8% lipids in the histological analysis at t=28d (P=0.0007). For vein grafts from ApoE*3-Leiden mice no differences in the lipid positive area was observed between the sPAI analysis or histological quantification. The percentages collagen present in the vein graft walls from both strains analyzed via sPAI and histological showed comparable results at t=28d. This indicates that sPAI was able to measure the amount of collagen and lipids in the vein graft wall.

## 4. Discussion

We have investigated the use of spectral photoacoustic imaging (sPAI) and superpixel photoacoustic unmixing (SPAX) method to non-invasively monitor different tissue distribution that can indicate vascular remodeling. Therefore, we herein studied the potential of photoacoustic image to automatically reveal functional and molecular parameters related vein graft tissue characteristics. The SPAX approach has been firstly tested via tissue-mimicking study and then validated in vivo murine model. Specifically, the in vivo longitudinal study included ApoE*3-Leiden and C57BL/6 groups at days 7, 14, and 28 after surgery. sPAI and SPAX data-driven analysis has enabled to automatically identify oxy/deoxy hemoglobin components and lipid and collagen. Histological analysis of collagen and lipids in the vein graft wall of ApoE*3-Leiden mice resulted in non-significant differences compared to the sPAI analysis.

The present study shows that componential analysis via photoacoustic imaging is an unbiased approach to discovering differences in composition within murine tissues. To our knowledge, this is the first time that a data-driven unmixing algorithm has been applied for unmixing spectral components as lipid and collagen in vivo within the NIR-I (680nm-970nm), where these tissue components are weaker absorbers. Thus, it is more challenging to detect in vivo lipid and collagen within NIR-I where hemoglobin is a prominent absorbers that obscure them. Were identified as melanin, oxidized hemoglobin, deoxidized hemoglobin, lipids, and collagen. These structural elements were identified within the vein grafts of mice in a 3D analysis.

Considering the current developments, PAI is a promising technology to follow the evolution of tissue remodeling in the carotid artery. The PAI technology can image functional/molecular information by using the optical absorption of endogenous tissue chromophores such as oxygenated and deoxygenated hemoglobin as well as lipids and collagen. Indeed, continuous assessments of the tissue molecular composition is essential to early identify the presence of plaques, as well as to follow up on the mechanical stability and rupture risk. Data analysis and algorithm development: Spectral photoacoustic imaging (sPAI), fused with linear unmixing method^19^, is a powerful modality used to generate distribution maps of the tissue chromophores by utilizing *a priori* information. Although this fitting-based approach yields acceptable results, it requires user interaction to provide the source spectral curves as an input. For translational research with patients, this type of supervised spectral unmixing can be challenging, as the spectral signature of the tissues differs with respect to the disease condition. Besides, light fluence attenuation along the imaging depth might induce spectral coloring that compromises the accurate unmixing and quantification of the chromophores. Thus, the linear unmixing algorithm can compromise the sensitivity and specificity of the imaging. An unsupervised spectral unmixing approach would be the ideal solution to overcome these limitations, and recently some studies have been reported, showing promising results^19-21^. A commonly used approach to identify tissue components is via linear unmixing and requires the input of expected tissue components. This is challenging because the known spectra could differ in disease models and limits the identification of novel components. To address this, we investigated the performance of a newly developed unsupervised superpixel photoacoustic unmixing (SPAX) framework^16^ that enables fully automatic spectral unmixing and accurate quantification of molecular components. This aims to reveal tissue biomarkers such as lipids and collagen, which are vital in the longitudinal monitoring of vascular remodeling in vivo and without bias.

In preclinical research, PAI has demonstrated its potential by visualizing tumor progression and recurrence. Clinically, PAI is used to monitor breast malignancies and lymph nodes in patients^22, 23^. Tumor oxygenation is a predictive risk factor for cancer aggressiveness^24^, which rely on the imaging of the vasculature. This imaging is performed in patients with tumors^25^ and patients with peripheral artery disease^26^.

Vein grafts are the preferred conduit to circumvent complex arterial occlusions, but are subjected to an accelerated form of atherosclerosis^18^. Lipids, which play a central role in the progression of these lesions, could serve as predictive markers for cardiovascular events. The spectral curve for lipids shows a distinct peak at 930 nm^27, 28^. Collagen, a connective tissue that provides elasticity and rigidity is associated with the stability of atherosclerotic lesions. Various isoforms of collagen exist and have been published ^27, 28,21^.

Lipids are the main driver of atherosclerotic plaque progression and follow-up could be a predictive risk factor for cardiovascular events. Collagen is the connective tissue that provided elasticity and rigidity to cope with arterial pressure. For in vivo PA imaging, the excitation light at the visible range or near-infrared I (NIR I: 650-900 nm) suffers from strong scattering in biological tissues, while the light in NIR II (NIRII: 1000-1700 nm) exhibits reduced light scattering. The identified collagen curve is comparable with the non-scatter collagen curve reported^29^.

This study demonstrates that componential analysis using photoacoustic imaging provides an unbiased approach to uncovering differences in composition within murine tissues. This methodology enhances our understanding of vein graft dynamics and holds promise for advancing the characterization of vascular diseases.

## Abbreviations

SPAX: superpixel photoacoustic unmixing
sPAI: photo acoustic imaging
BAT: brown adipose tissue
WAT: white adipose tissue

## Supplemental figures

**Figure S1.**
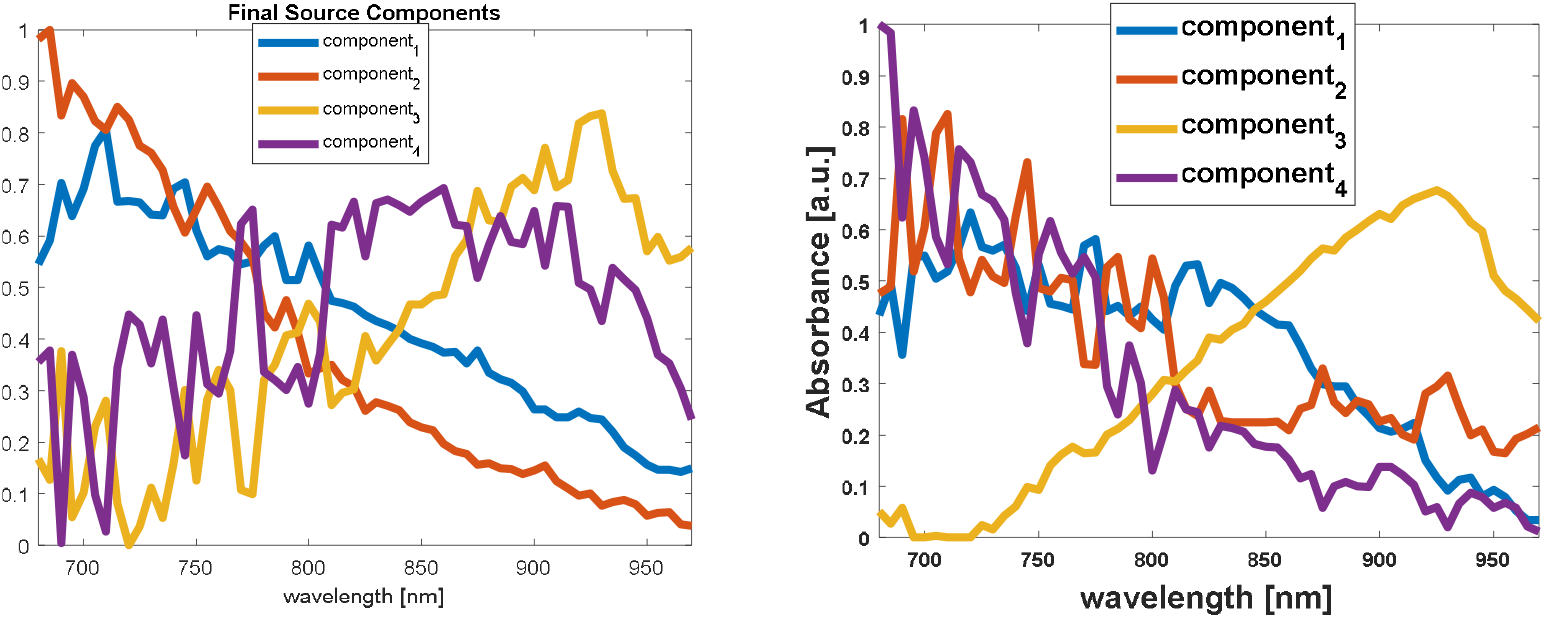
windowing to enhance spectral curve discremination. Example spectral curve before windowing (A) and windowing applied (B).

